# Novel antimicrobial peptide discovery using machine learning and biophysical selection of minimal bacteriocin domains

**DOI:** 10.1101/314740

**Authors:** Francisco R. Fields, Stephan D. Freed, Katelyn E. Carothers, Md Nafiz Hamid, Daniel E. Hammers, Jessica N. Ross, Veronica R. Kalwajtys, Alejandro J. Gonzalez, Andrew D. Hildreth, Iddo Friedberg, Shaun W. Lee

## Abstract

Bacteriocins are ribosomally produced antimicrobial peptides that represent an untapped source of promising antibiotic alternatives. However, inherent challenges in isolation and identification of natural bacteriocins in substantial yield have limited their potential use as viable antimicrobial compounds. In this study, we have developed an overall pipeline for bacteriocin-derived compound design and testing that combines sequence-free prediction of bacteriocins using a machine-learning algorithm and a simple biophysical trait filter to generate minimal 20 amino acid peptide candidates that can be readily synthesized and evaluated for activity. We generated 28,895 total 20-mer peptides and scored them for charge, α-helicity, and hydrophobic moment, allowing us to identify putative peptide sequences with the highest potential for interaction and activity against bacterial membranes. Of those, we selected sixteen sequences for synthesis and further study, and evaluated their antimicrobial, cytotoxicity, and hemolytic activities. We show that bacteriocin-based peptides with the overall highest scores for our biophysical parameters exhibited significant antimicrobial activity against *E. coli* and *P. aeruginosa*. Our combined method incorporates machine learning and biophysical-based minimal region determination, to create an original approach to rapidly discover novel bacteriocin candidates amenable to rapid synthesis and evaluation for therapeutic use.

## Introduction

Many bacteria have become resistant to conventional antibiotics, necessitating the discovery of novel antimicrobial compounds^1^. However, pharmaceutical antibiotic development has declined chiefly due to brief usability window of existing antibiotic scaffolds ^2^. To combat the lack of novel antimicrobial discovery, many bioinformatic approaches have been developed to mine the genomes of bacteria for natural products^3^. One promising class of natural products are bacteriocins, the ribosomally produced antimicrobial peptides of bacteria^4,5^. These chemically and functionally diverse peptides are divided into two main classes. The class I bacteriocins include extensive post-translational modifications in their final form; for example, nisin is a commonly studied bacteriocin whose features include post-translational modifications such as lanthionine and methyllanthionine^6^. Enterocin AS-48, another class I bacteriocin, undergoes head-to-tail circularization^7^. The class II bacteriocins primarily consist of peptides that do not undergo post-translational modification^4^. Bacteriocins are often located in genetic clusters containing the structural gene encoding the precursor peptide, as well as the context genes necessary for bacteriocin maturation, export, and immunity. The bacteriocin structural gene is often expressed as a prepropeptide, consisting of the unmodified bacteriocin functional domain and an N-terminal leader sequence. Upon installation of the post-translational modifications and cleavage of the leader peptide, the mature peptide is exported by an ABC-cassette type transporter^4,5,8^.

Genome mining approaches have taken advantage of the bacteriocin operon-like structure to identify novel bacteriocin candidates through two approaches: identification of the bacteriocin precursor gene or identification of bacteriocin context genes^3, 8–10^. Online genome mining tools, such as BAGEL, and bacteriocin databases, such as BACTIBASE, allow investigators to identify and classify putative bacteriocins based on their homology to other known bacteriocin genes^9,10^. Similar tools, such as anti-SMASH, have been expanded to not only identify putative bacteriocins, but also secondary metabolites and other genetically identifiable antibiotics^3,8,11,12^.

Large sequence heterogeneity and a small number of experimentally determined bacteriocins, as well as the small size of most structural genes (30-150aa) have presented challenges in identifying novel bacteriocins using BLAST and other sequence similarity approaches ^8^. To overcome these problems, some bacteriocin prediction software identify novel bacteriocins by searching for conserved context genes of the bacteriocin operon^8,13^. The <u>b</u>acteriocin <u>o</u>peron and gene block <u>a</u>ssociator (BOA) identifies context genes through homology-based genome searches^8^. BOA has identified 95% of BAGEL annotated bacteriocins in addition to 1,033 putative bacteriocins not identified by BAGEL. ClusterFinder, another context gene based approach, has been used to mine the genomes of human commensal organisms. This approach led to the identification of the novel thiopeptide bacteriocin lactocillin^13^. Another tool, MetaRiPPquest, connects genomic bacteriocin predictions to tandem mass spectrometry data. Peptidogenomic approaches attempt to bridge the gap between computational and *in vitro* identification^14^. While context-based approaches seem to circumvent the need for sequence similarity, novel methods that move away from homology-based mining tools are still needed. Recently, *k*-mer based machine learning approaches have been used successfully to classify protein sequences without the need for homology^15,16^.

Regardless of the genome mining approach, *in vitro* verification of the antimicrobial activity of computationally identified putative bacteriocins also remains a major challenge due to several factors. First, bacteriocins have diverse mechanisms of action with most having specific targets within or on host cells. This is especially true for the class I bacteriocins; for example, microcin B17 (MccB17) inhibits the activity of DNA gyrase while nisin inhibits peptidoglycan synthesis by binding to lipid II^17–19^. Even class II bacteriocins can have extremely specific targets; for example, lactococcin A targets the mannose phosphotransferase system to induce pore formation^5,20^. Secondly, bacteriocins may exhibit a very narrow spectrum of activity and be highly specific against a competitor strain. Many bacteriocins produced by lactic acid bacteria will only kill other closely related species such as *Lactobacillus*, *Enterococcus*, and *Listeria^5^*. Finally, some bacteriocins may not have bacterial targets. Streptolysin S (SLS) is structurally similar to MccB17 as both are thiazole-oxazole modified microcins; however, SLS is a virulence factor that promotes invasion upon Group A *Streptococcus* infection^21,22^. Other bacteriocin-like peptides may act as signaling molecules, as nisin and subtilosin have both been implicated as autoregulators acting as autocrine signaling peptides at distinct concentrations ^11,19^. However, some bacteriocins target the bacterial membrane in a non-specific fashion through electrostatic and hydrophobic interactions, including enterocin AS-48 and sakacin^5,7,23^.

Recently, we have shown that membrane targeting bacteriocins can serve as templates for the efficient design of synthetic antimicrobial peptides^24^. Using the AS-48 homologue, safencin AS-48, we created a synthetic peptide corresponding to the membrane interacting region of enterocin AS-48^7,25^. This region is a cationic, hydrophobic, alpha-helical peptide that abstracts the full-length 70 amino acid bacteriocin to a 25 amino acid peptide. Interestingly, these biophysical qualities are very similar to synthetic antimicrobial peptides derived from eukaryotic sources, whose activity relies on their overall positive charge, conformation, and amphipathicity^26–29^. To determine if these biophysical guidelines of antimicrobial peptide design could select for regions of putative membrane-targeting bacteriocins, we wrote a script to scan for 20 amino acid stretches at a time along the length of a putative bacteriocin and score each 20-mer for charge, alpha-helical propensity, and hydrophobic moment. Upon chemical synthesis and antimicrobial testing of a set of these 20-mers, we observed that peptide candidates with the highest scores in all three categories exhibited significant antimicrobial activity. This approach represents a method by which membrane-targeting regions of putative bacteriocins can be rapidly selected, synthesized, and verified *in vitro*. We propose that peptides discovered through this process could then serve as scaffolds for subsequent optimization and eventual therapeutic development.

## Materials and Methods

### Initial selection of candidate bacteriocins

We selected an initial set of putative novel bacteriocins using a word embedding algorithm, Word2vec, as described previously^16^. Briefly, we created a vocabulary of all possible 8,000 amino-acid trimers. Each trimer is then represented as a vector, which captures the probabilities of that trimer being in the neighborhood of other trimers, also known as the skip-gram model^15^. Each protein sequence was then represented as the sum of vectors representing the trimers comprising the protein. We then trained several supervised learning models with a positive set of 346 known bacteriocins and a negative set of the same size. The best performing method, and support vector machine (SVM) was then used to discover the set of 676 putative bacteriocins used in this study. In essence, the machine learning algorithm employed thus generates a list of new bacteriocin-like sequences that preserve key evolved features of natural bacteriocins products. We eliminated from the list all known bacteriocins which were discovered using BLAST against GenBank with an e-value of 10^-3^ or less, and which were annotated as bacteriocins. The result was a set of 676 putative bacteriocins, not obviously homologous, by sequence similarity, to existing bacteriocins.

### Biophysical selection of 20-mer peptides

Using a sliding window, we generated 28,895 20-mers from the 676 predicted peptides, and calculated the following biophysical parameters for each 20-mer candidate: (1)Charge, (2) Helicity, and (3) Hydrophobic moment (Figure 1)^30–32^. Net charge was calculated as a sum of the charge for each amino acid at pH 7. Helicity was calculated as a sum of the Chou-Fasman probabilities of each amino acid. Finally, the hydrophobic moment was calculated using the hydrophobicity values for each residue assuming that the 20-mer peptides would adopt an alpha helical structure. For each of the biophysical parameters, the 20-mers were ranked as high, middle, or low based on the range of scores within that parameter (Figure 2A).

**Figure 1:**
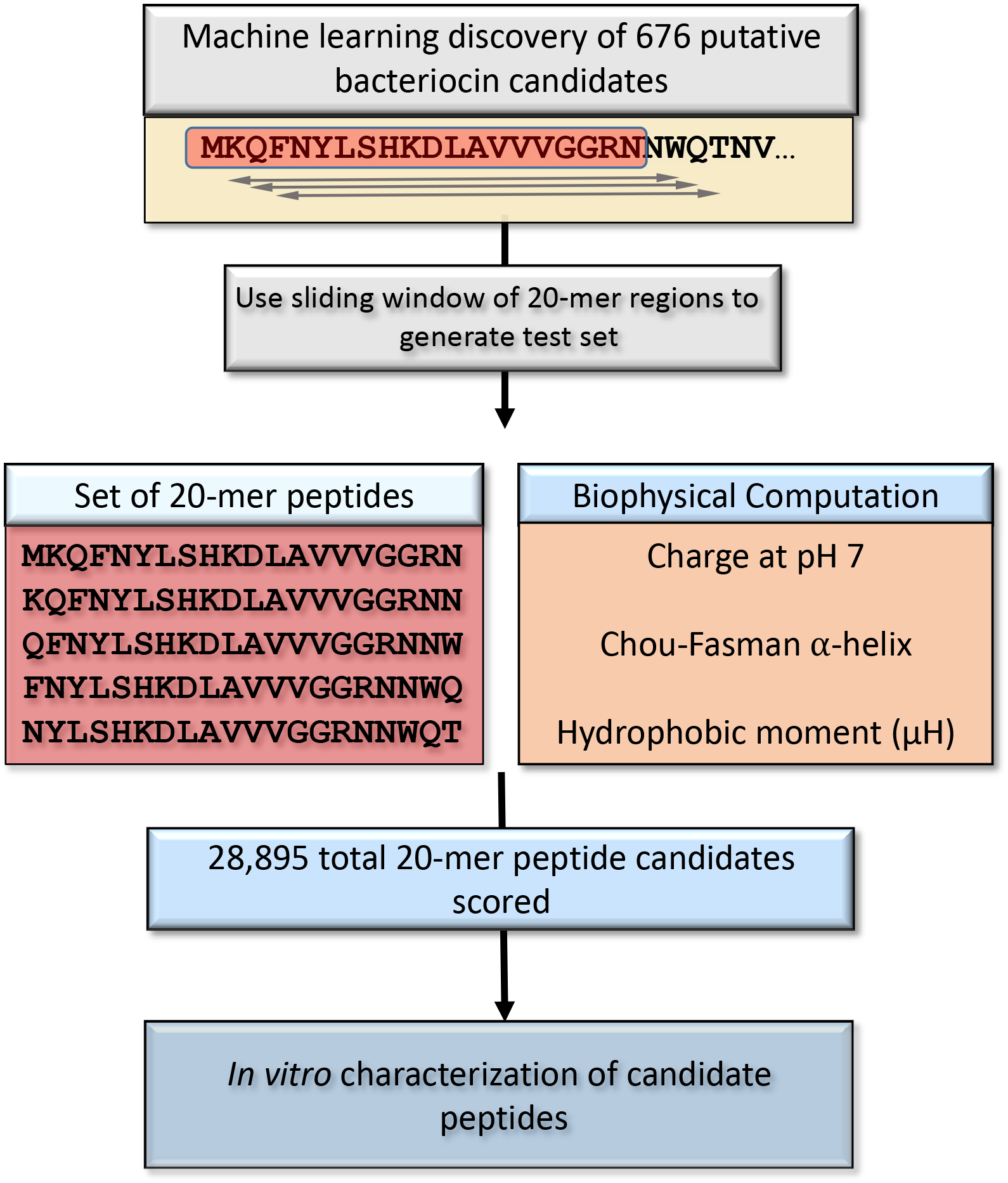
Overall strategy for selection of bacteriocins for synthesis. Machine learning set of 676 putative bacteriocins was used to generate overlapping 20-mer peptide candidates. 28,895 20-mers were scored and ranked for charge, helicity, and hydrophobic moment. A representative sample of 16 peptides were selected for synthesis and in vitro characterization in this study.

**Figure 2:**
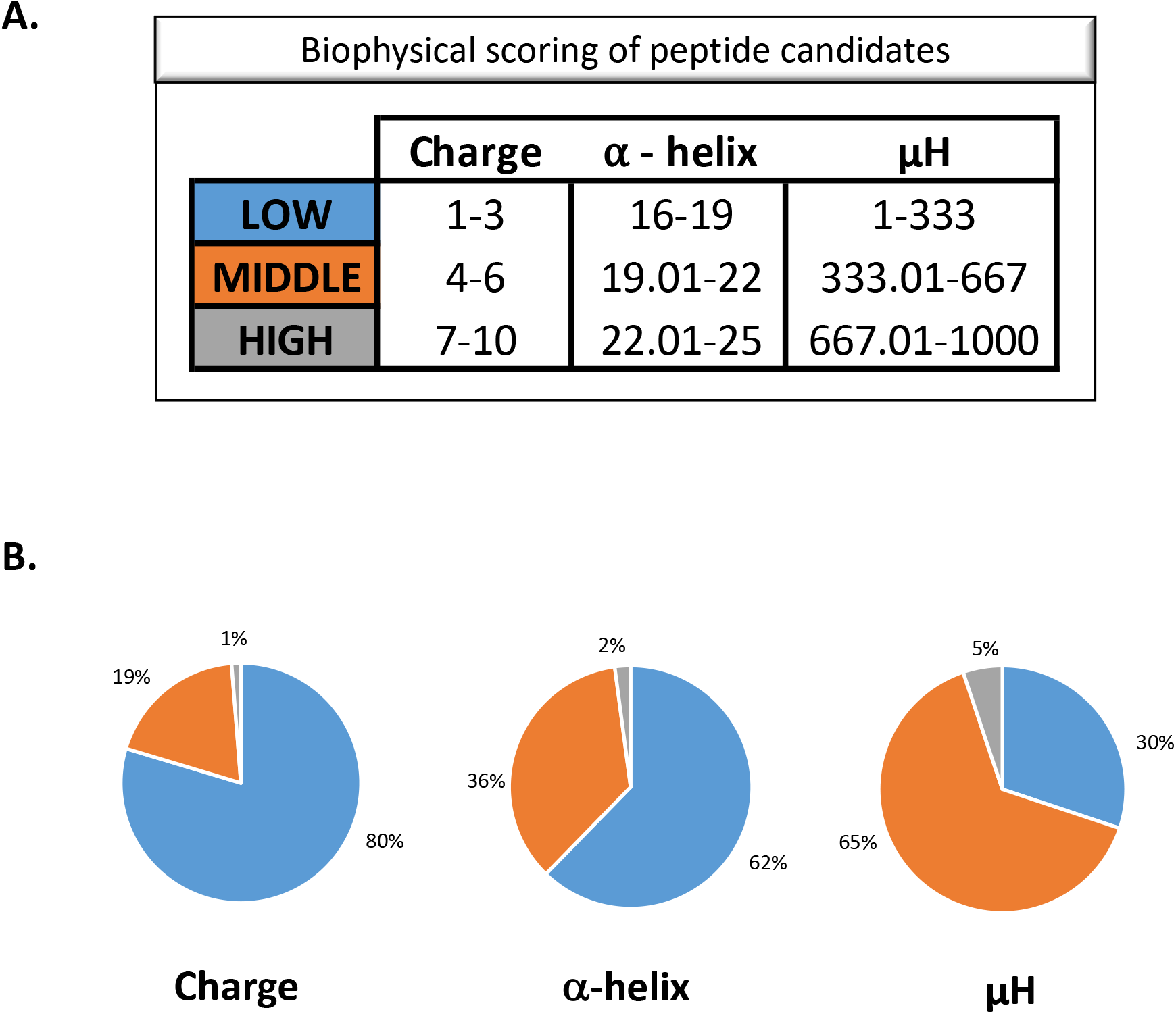
Scoring breakdown of biophysical computational parameters of the candidate peptides. **A**. Peptides were divided into high (grey), middle (orange), and low (blue) groups based on their charge, helicity, and hydrophobic moment scores. **B**. Most of the peptides scored low to middle with only a small percentage scoring high for each of the biophysical parameters.

### Peptide Selection and Synthesis

To evaluate our biophysical parameter scores, we selected a total of sixteen 20-mer peptides for synthesis and further experimentation. Peptides were synthesized by Genscript (Piscataway, NJ), to >95% purity and verified by HPLC and mass spectrometry. All peptides were dissolved in DMSO for subsequent experimentation (Thermo Fischer).

### Bacteria and Growth Conditions

*E. coli* BL-21 (Thermo Fischer) and *P. aeruginosa* PAO1 (gift from J. Shrout at the University of Notre Dame) were grown in LB broth Miller (EMD chemicals, Gibbstown NJ). *Staphylococcus aureus* USA300 was grown in Todd Hewitt broth (Neogen Corporation, Lansing, MI). All cultures were grown at 37 °C.

### Antimicrobial Activity Assays

Minimal inhibitory concentrations (MICs) of the 20-mer peptides were determined via microtiter dilution assay^33^. Briefly, dilute bacterial cultures were added to a series of serial two-fold dilutions of peptide in Mueller-Hinton broth (Thermo Fischer). The lowest concentration at which no bacterial growth was observed after overnight incubation at 37 °C was defined as the MIC. If an MIC could be determined, cultures from the MIC experiment were plated and incubated overnight at 37 °C. The concentration at which no colonies were visible after overnight incubation was defined as the minimal bactericidal concentration (MBC).

### Antibiofilm Formation Assays

Antibiofilm activity of the peptides were assessed using USA 300 and PAO1. For USA300 biofilms, overnight cultures grown in TSB (Sigma-Aldrich) were diluted 1:100 in TSB .1% glucose 1% NaCl with or without peptide^34^. For PAO1 biofilms, overnight cultures grown in LB were diluted 1:100 in M63 1mM MgSO4 and .4% arginine with or without peptide^35^. Samples were incubated for 24 hours in a microplate. Planktonic cells were removed from the wells and the biofilms were washed three times with ddH2O. Biofilms were then stained with .1% crystal violet, washed three times with ddH2O, and resuspended in 30% acetic acid36. These were then quantified by OD 550 reading on an Synergy Microplate Reader (Biotek).

### Peptide Cytotoxicity Assays

Eukaryotic cytotoxicity was determined by ethidium homodimer and hemolysis assays. Ethidium homodimer assays were carried out with HaCaT cells in 24 well culture dishes grown to 90% confluency. Medium was aspirated, and cells were washed with PBS (Thermo Fischer). Peptide in fresh DMEM (Dibco) was added to the cells at the desired concentration. Cells were incubated with peptide for 16 hours. Medium was aspirated, and cells were washed with PBS. Cells were incubated in 4 μM ethidium homodimer (Molecular Probes) in PBS for 30 minutes. Fluorescence was determined by 528 excitation and 617 nm emission and a cutoff value of 590 nm. Saponin (.1%) was then added to each well and incubated for 20 minutes. The fluorescence was read again. Percent membrane permeabilization was determined by dividing the initial fluorescence by the second fluorescence reading. For hemolysis assays, 100 μL of sheep red blood cells (RBCs) were washed 3 times in cold PBS. Washed cells were resuspended in 25 ml of PBS. Triton, PBS, or peptide in 10% DMSO/PBS were added to 180 μL of resuspended RBCs and incubated at 37°C for 1 hour. Samples were read at 450 nm. Data was expressed as percent hemolysis by relativizing to the Triton and PBS controls.

## Results

### Design and biophysical selection of 20-mer minimal bacteriocins

From the initial set of 676 putative novel bacteriocins using the word embedding algorithm, Word2vec, 28,895 total 20-mer bacteriocin peptide candidates were generated (Figure 1). Each peptide was then assigned a low, middle, or high ranking for each of the biophysical parameters based on the range of scores within that parameter (Figure 2A). For example, a peptide with a net charge of 5, a helical score of 17, and a μH of 900 would rank middle for charge, low for helicity, and high for μH (Figure 2A).

80% of the 20-mers received a low ranking for charge (a net positive charge between +1 and +3) while only 1% ranked high (Figure 2B). For the hydrophobic moment values, a majority of the peptides also ranked low (any hydrophobic moment value below 333) with only 5% receiving a high score (Figure 2B). However, for the helicity score, a majority of the peptides, 65%, fell into the middle range of scores between 19 and 22 with only 2% scoring high for helicity (Figure 2B). It is important to note that the hydrophobic moment and helicity scores may not truly represent these parameters for the peptides as the propensity to form a beta sheet was not taken into consideration when calculating these values.

### Peptide Selection for Chemical Synthesis

Many cationic antimicrobial peptides will adopt an amphipathic alpha helical conformation. Therefore, we reasoned that of the peptides generated by our script those ranking high in all three biophysical categories would yield the most antimicrobial activity. Of the sixteen peptides selected for synthesis, peptides 1 and 2 ranked low for all three biophysical parameters while peptides 3 and 4 ranked high for the three parameters (Table 1). The remaining 12 peptides were randomly selected from all 20-mers ranking middle in at least one category and high for the remaining parameters (Table 1).

**Table 1:**
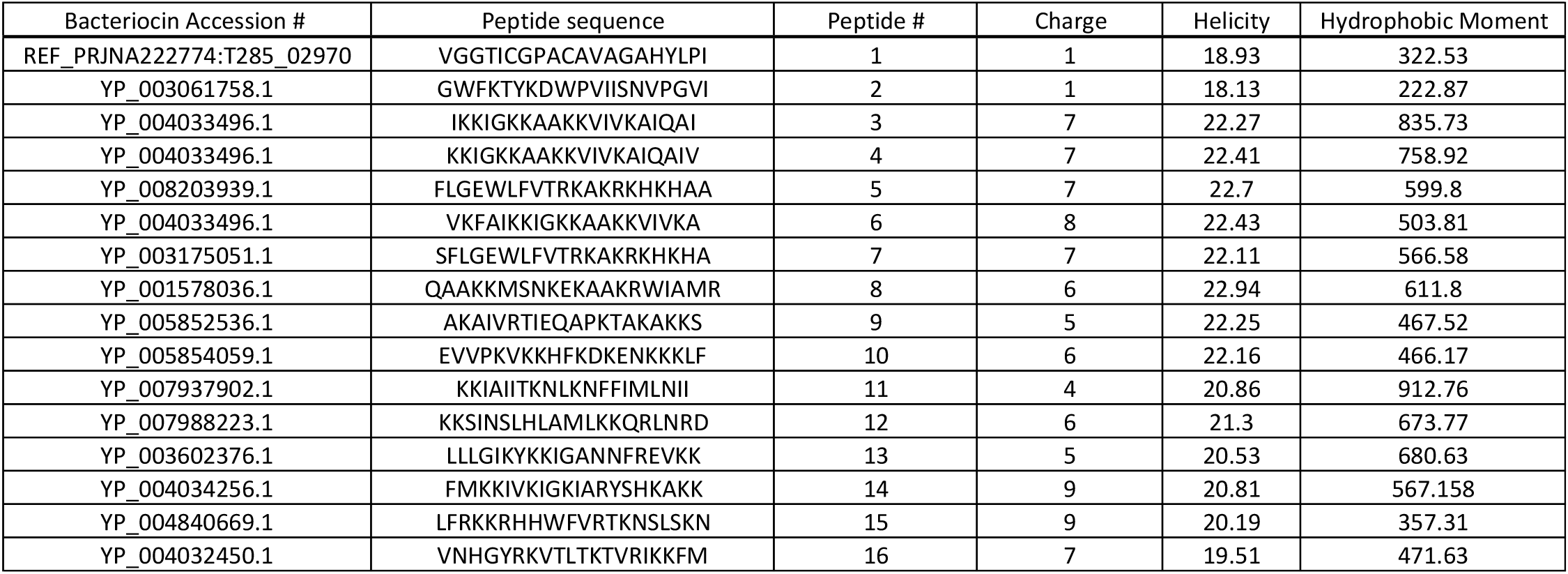
20-mers selected for synthesis and their corresponding biophysical scores.

### PEP-FOLD prediction of secondary structure

To determine if our biophysical selection criteria were able to accurately predict an amphipathic alpha helical structure of the peptides selected for synthesis, we modeled their secondary structure using the PEP-FOLD online tool. For peptides 1 and 2, which received low scores for helicity and hydrophobic moment, their structures are predicted to exist as a majority random coil (Figure 3A). In contrast, peptides 3 and 4, having high scores for helicity and hydrophobic moment, are predicted to exist as fully extended alpha helices with clear clustering of the polar and charged amino acids to one side of the helix and the hydrophobic residues on the other, indicative of a strong hydrophobic moment (Figure 3B). Peptides 5 through 10 have a high helicity score; however, the modeling predictions expect unstructured regions owing to helix-breakering residues glycine and proline that occur within their sequences (Figure 3 and Table 1). All of these peptides also received middle scores for their hydrophobic moment which is visible as hydrophobic residues within the polar face of the helix, such as peptide 6, and charged amino acids within the hydrophobic face, such as peptide 9. Interestingly, peptide 11 is predicted to exist as a beta sheet (Figure 3E). The biophysical calculator only takes into account the Chou-Fasman residue helical propensity score and does not calculate the individual likelihood of forming a beta sheet; therefore, peptides with a higher sheet propensity were not excluded from the list of peptides for synthesis. Finally, the rest of the peptides are predicted to adopt various helical structures with differing amphipathic characteristics (Figure 3).

**Figure 3:**
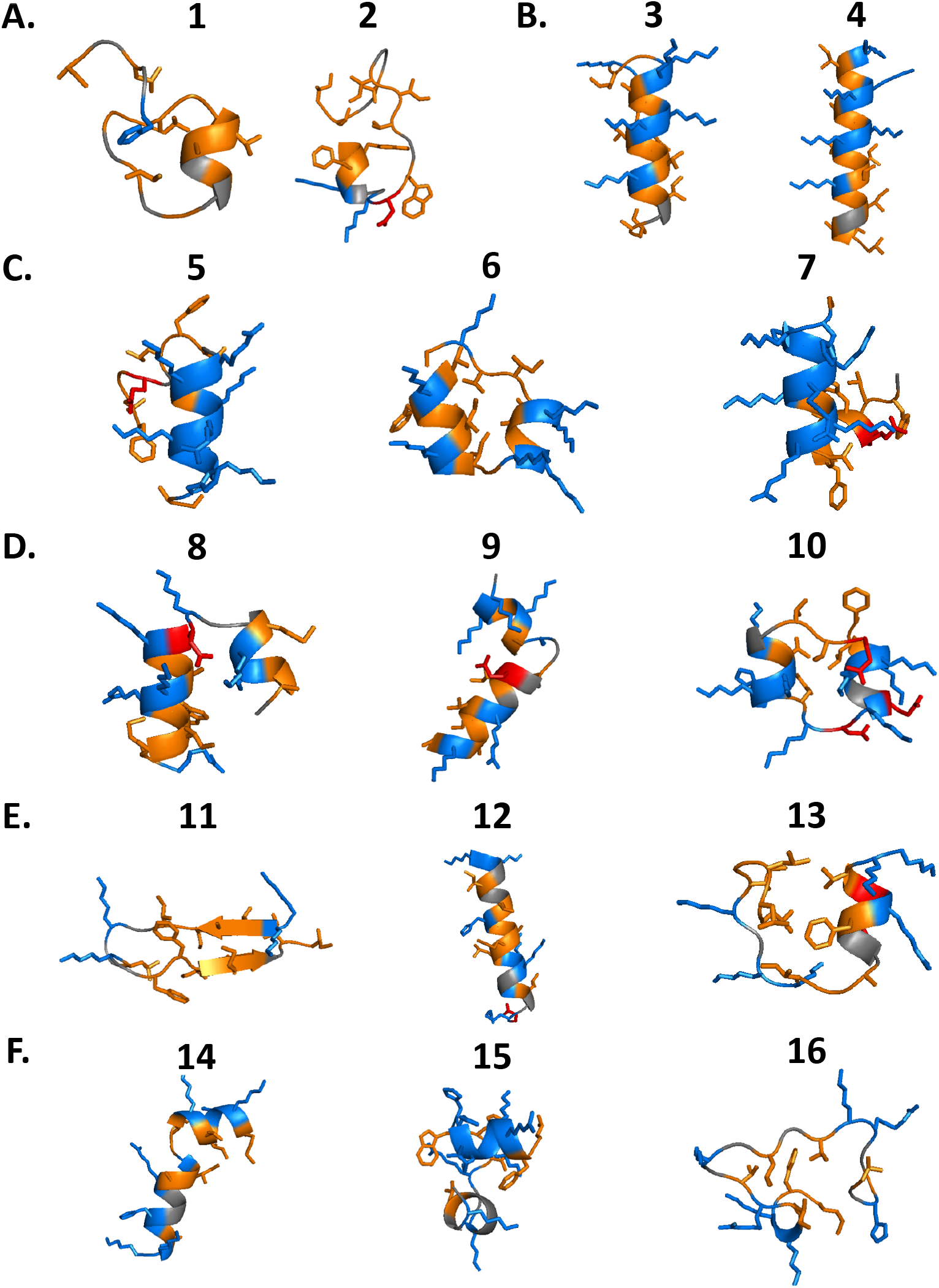
PEP-FOLD models of the peptides selected for synthesis **A.** peptides 1 and 2, **B.** 3 and 4, **C.** 5,6, and 7, **D.** 8, 9, and 10, **E.** 11, 12, and 13, and **F.** 14, 15, and 16. Basic, acidic, and hydrophobic residues are in blue, red, and orange respectively.

### Antimicrobial Properties of Synthetic 20-mers

The peptides were assessed for their minimal inhibitory concentration (MIC) and minimal bactericidal concentration (MBC) on *Escherichia coli, Staphylococcus aureus,* and *Pseudomonas aeruginosa* (Table 2). As expected, peptides 1 and 2, which scored low in all three biophysical parameters, did not have activity against any of the organisms tested. Peptides 3 and 4, which scored high in all three biophysical parameters, exhibited antimicrobial activity against both *E. coli* and *P. aeruginosa* (Table 2). Peptides 5, 6, and 7 scored high in charge and helicity and middle in hydrophobic moment (Table 1). Interestingly, these peptides showed a range of antimicrobial activities (Table 2). Peptide 6 was more efficient at inhibiting the growth of *P. aeruginosa* (MIC = 32 μM) than *E. coli* (MIC =128 μM). Peptides 5 and 7 were much less active than peptide 6 despite having similar values for their biophysical scores (Tables 1 and 2). Peptides 8, 9, and 10 scored high for helicity with middle scores for charge and hydrophobic moment. These peptides did not have any antimicrobial activity against the organisms tested. This overall trend continued for the rest of the peptides tested. Indeed, peptides scoring high in any one of the biophysical parameters with only middle scores for the others (peptides 8-16) did not have any antimicrobial activity. We did not test any of the peptide candidates at concentrations above 128μM, so biological activities at higher concentrations cannot be ruled out.

**Table 2:** MICs and MBCs of the synthetic 20-mer bacteriocins against *S. aureus*, *E. coli*, and *P. aeruginosa.*

| Peptide # | MICs Table |  |  |  |  |  |
| --- | --- | --- | --- | --- | --- | --- |
|  | *Pseudomonas aeruginosa* |  | *Escherichia coli* |  | *Staphylococcus aureus* |  |
|  | MIC (μM) | MBC (μM) | MIC (μM) | MBC (μM) | MIC (μM) | MBC (μM) |
| 1 | > 128 | > 128 | > 128 | > 128 | > 128 | > 128 |
| 2 | > 128 | > 128 | > 128 | > 128 | > 128 | > 128 |
| 3 | 64 | > 128 | 32 | 64 | > 128 | > 128 |
| 4 | 16 | 32 | 2 | 4 | > 128 | > 128 |
| 5 | > 128 | > 128 | 128 | > 128 | > 128 | > 128 |
| 6 | 32 | 128 | 128 | > 128 | > 128 | > 128 |
| 7 | 128 | > 128 | 128 | > 128 | > 128 | > 128 |
| 8 | > 128 | > 128 | > 128 | > 128 | > 128 | > 128 |
| 9 | > 128 | > 128 | > 128 | > 128 | > 128 | > 128 |
| 10 | > 128 | > 128 | > 128 | > 128 | > 128 | > 128 |
| 11 | > 128 | > 128 | > 128 | > 128 | > 128 | > 128 |
| 12 | > 128 | > 128 | > 128 | > 128 | > 128 | > 128 |
| 13 | > 128 | > 128 | > 128 | > 128 | > 128 | > 128 |
| 14 | > 128 | > 128 | 128 | > 128 | > 128 | > 128 |
| 15 | > 128 | > 128 | > 128 | > 128 | > 128 | > 128 |
| 16 | > 128 | > 128 | > 128 | > 128 | > 128 | > 128 |

### Inhibition of Biofilm Formation by the Synthetic 20-mers

Despite not having a true MIC, we observed that peptides 11 and 16 were able to significantly reduce the overnight growth of *S. aureus* cultures (Supplementary Figure 1A-B). To investigate if these peptides were exerting antibiofilm effects, we employed the biofilm formation assay. Upon incubation with peptide 11 for 24 hours in biofilm inducing media, we observed a significant decrease in USA 300 biofilm formation down to a concentration of 4 μM (Supplementary Figure 1C). This trend was also observed for peptide 16; however, this only inhibited biofilm formation down to 16 μM (Supplementary Figure 1D) Finally, to determine if these peptides could inhibit the biofilms of other bacteria we used *P. aeruginosa*. Peptides 11 and 16 exhibited no bacteriostatic effects on PAO1 (Supplementary Figure 2A-B). However, these peptides exerted mild antibiofilm formation activity down to 16 μM (Supplementary Figure 2C-D). In addition to identifying peptides with potent antimicrobial activity, we have also identified peptides with antibiofilm activity.

### Peptide mammalian cell cytotoxicity

To determine if our biophysical parameters were able to select for peptides with affinity for bacterial membranes instead of mammalian membranes, we assessed their ability to compromise the membranes of erythrocytes and keratinocytes. Fourteen of the peptides exhibited no hemolytic activity even at high concentrations (Supplementary Table 1). However, peptides 2 and 10 exhibited increased levels of hemolysis at only the highest concentrations (128 μM). Cytotoxicity to keratinocytes was interrogated using the ethidium homodimer assay. We observed that all of the peptides were unable to cause cell death when incubated with HaCaT cells for 16 hours at the highest concentrations. Together, these data indicate that these peptides generally do not target mammalian membranes.

## Discussion

Bacteriocins are a barely-tapped source of highly diverse antimicrobials. However, verifying the antimicrobial activity of putative bacteriocins can be difficult due to the potentially narrow activity spectra and highly diverse mechanisms^4,37^. Additionally, traditional methods of natural bacteriocin isolation as well as heterologous expression strategies are complicated by purification limitations and low yield^38–40^. Here we describe a complete strategy by which *de novo* mining of bacteriocins can be parsed using a biophysical algorithm to identify minimally active bacteriocin peptide candidates. Biophysical selection was done by focusing on three parameters that have been implicated in the activity of membrane active antimicrobial peptides: helicity, charge, and amphipathicity^30–32,41–43^. Our strategy for employing predictive algorithms with biophysical selection and minimal domain candidate design allows for the development of completely novel, highly active, synthetic bacteriocins that have wide applicability as antimicrobial compounds. Previous studies have shown that synthetic peptide variants of full length bacteriocins can be used to approximate their antimicrobial function. For example, linear variants of enterocin AS-48, a circular bacteriocin consisting of five alpha helices, have been shown to retain some of the antimicrobial activity of the parent^44,45^. The antimicrobial action was shown to be dependent upon the cationic and hydrophobic residues present within helices four and five that are designated as the membrane-interacting region^25^. Recently, we published a strategy whereby the membrane-interacting region in an AS-48 homologue was used as a template to create a series of small, optimized antimicrobial peptides^24^. This establishes a precedent by which synthetic peptides can be used to approximate the activity of the full length bacteriocin. We have built upon these previous studies by utilizing the biophysical parameters of synthetic antimicrobial peptide design to select for membrane interacting regions of putative bacteriocins^25^. We observed that peptides with the highest scores for the biophysical parameters of charge, helicity, and hydrophobic moment were the most active against the bacteria tested (Table 2). Interestingly, the only two peptides to meet these criteria were from the same putative bacteriocin. It is therefore highly likely that this putative bacteriocin works in a membrane active manner ^29,42,43^. The interpretation of these data becomes confounded for the peptides whose biophysical parameters begin to receive middle scores. For example, peptide 6, with a middle score for hydrophobic moment, is a more effective antimicrobial against *P. aeruginosa*, MIC = 32, than *E. coli*, MIC = 128. This observation is in contrast to the activities of the high scoring peptides, 3 and 4, whose antimicrobial activities were higher against *E. coli*. Therefore, it may be possible to tune antimicrobial *specificity* by modifying the biophysical scores^46,47^. While most research has focused on modification of these parameters and their effects on eukaryotic cytotoxicity and overall antimicrobial activity few have examined how these parameters tune the specificity of these compounds to specific bacteria^48,49^.

There are some drawbacks to this approach. While it seems that our approach has selected for antimicrobial regions of putative bacteriocins, it is also possible that using a minimal synthetic peptide strategy has decoupled the function of the synthetic bacteriocin from the function of the full sequence. Enterocin AS-48 undergoes dimer formation and then subsequent tertiary structural changes before inserting itself into the membrane of target bacteria^50^. However, synthetic AS-48 peptides lose this ability to dimerize and work in a mechanism more akin to carpet or pore models of synthetic antimicrobial peptide activity^25^. Therefore, some of the antimicrobial function and specificity inherent in bacteriocins will be lost by utilizing synthetic minimal versions. Finally, our approach cannot verify the activity of bacteriocins which do not target the bacterial membrane or whose biophysical characteristics change upon post-translational modification^4,5,11^.

Despite these drawbacks, the techniques described herein have potential for linking *de novo* computational bacteriocin discovery with immediate therapeutic development. With the increasing amount of computational work being done to predict novel antimicrobial compounds there is a mounting need to verify their antimicrobial activity *in vitro^8–10^*. Our method validates the use of machine learning algorithms to further mine genomic information for potential bacteriocins candidates that can be refined using biophysical scripting parameters and size optimization for rapid synthesis and testing. The lack of mammalian cell cytotoxicity in our synthesized peptide set indicates that selecting minimal bacteriocin candidates based on the specific set of biophysical parameters that we have established will select for candidates that specifically target bacterial membranes, a highly valuable outcome from our studies (Table 2 and Supplementary Table 1). Many current synthetic antimicrobial peptides used to treat human disease have been built around an existing scaffold from eukaryotes^51^. Omiganan, derived from magainin of the African three-toed frog, is currently being developed as a topical antimicrobial for the treatment of diabetic foot ulcers^52^. In contrast, relatively few bacteriocins have been developed for the treatment of disease^51,53,54^. Our strategy to combine machine learning algorithms for *de novo* bacteriocin discovery along with biophysical refinement and minimal design represent a particularly robust workflow for the development of new antibiotic compounds. These synthetic bacteriocin scaffolds could be further refined via iterative testing and data collection for efficacy and selectivity.

## Acknowledgements

IF was funded, in part, by NSF awards ABI-1551363 and ABI-1458359. SL was funded by NIH Innovator Award 1DP2 OD008468-01 and an Eck Institute for Global Health Pilot Project Program. FF is supported by an NSF-GRFP fellowship and a National GEM Consortium Fellowship.

---

**Supplemental Figure 1:**
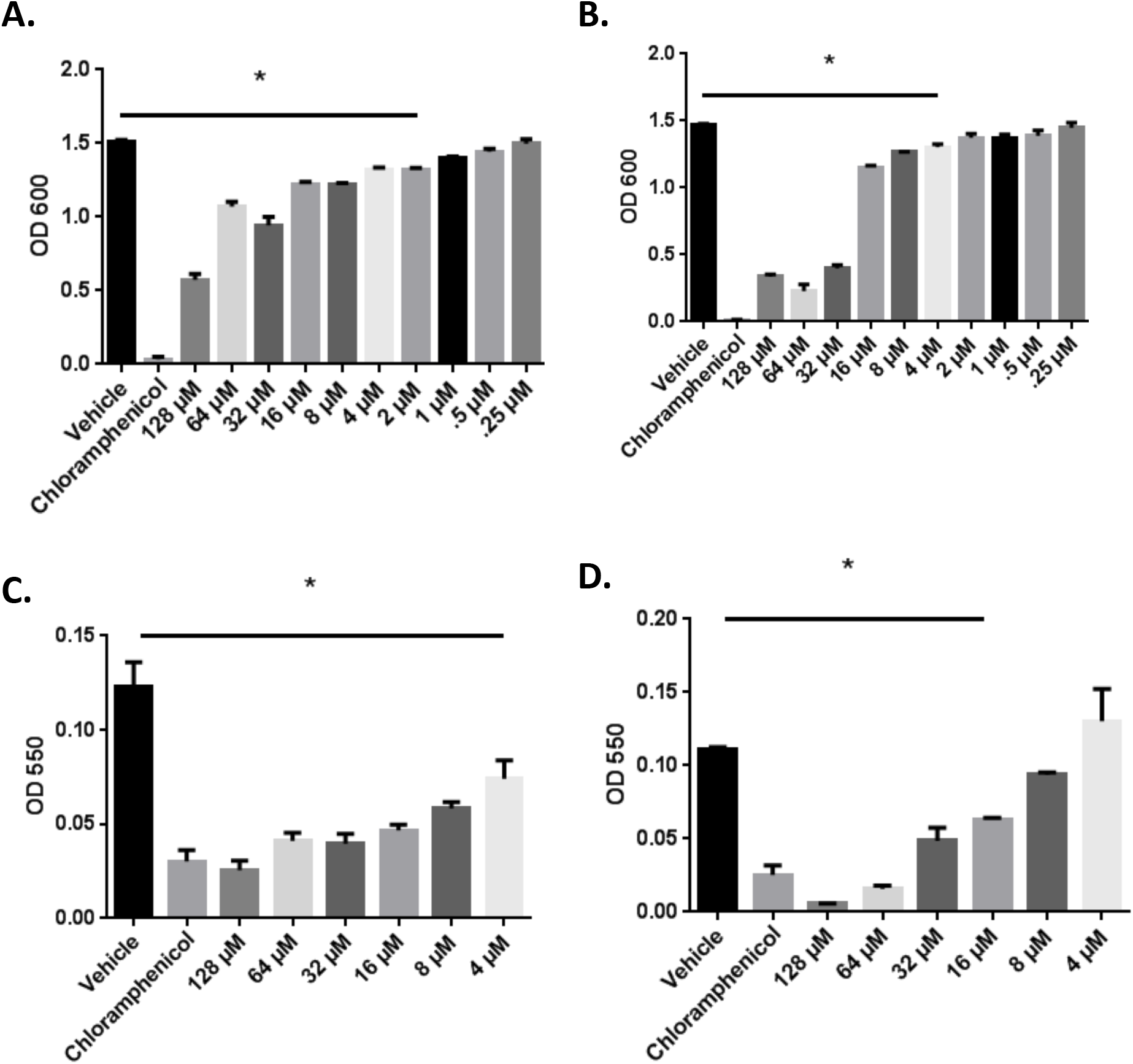
Antibiofilm activities of peptides 11 and 16 on *S. aureus*. **A.** Peptide 11 exhibits a bacteriostatic effect and **B.** peptide 16 exhibits a bacteriostatic effect **C.** Peptide 11 inhibits biofilm formation at all concentrations tested. **D.** Peptide 16 inhibits biofilm formation to 16 μM. Data is representative of 3 biological replicates. P-values were determined via one-way ANOVA. A * indicates a significant difference determined via Tukey HSD compared to the vehicle control.

**Supplemental Figure 2:**
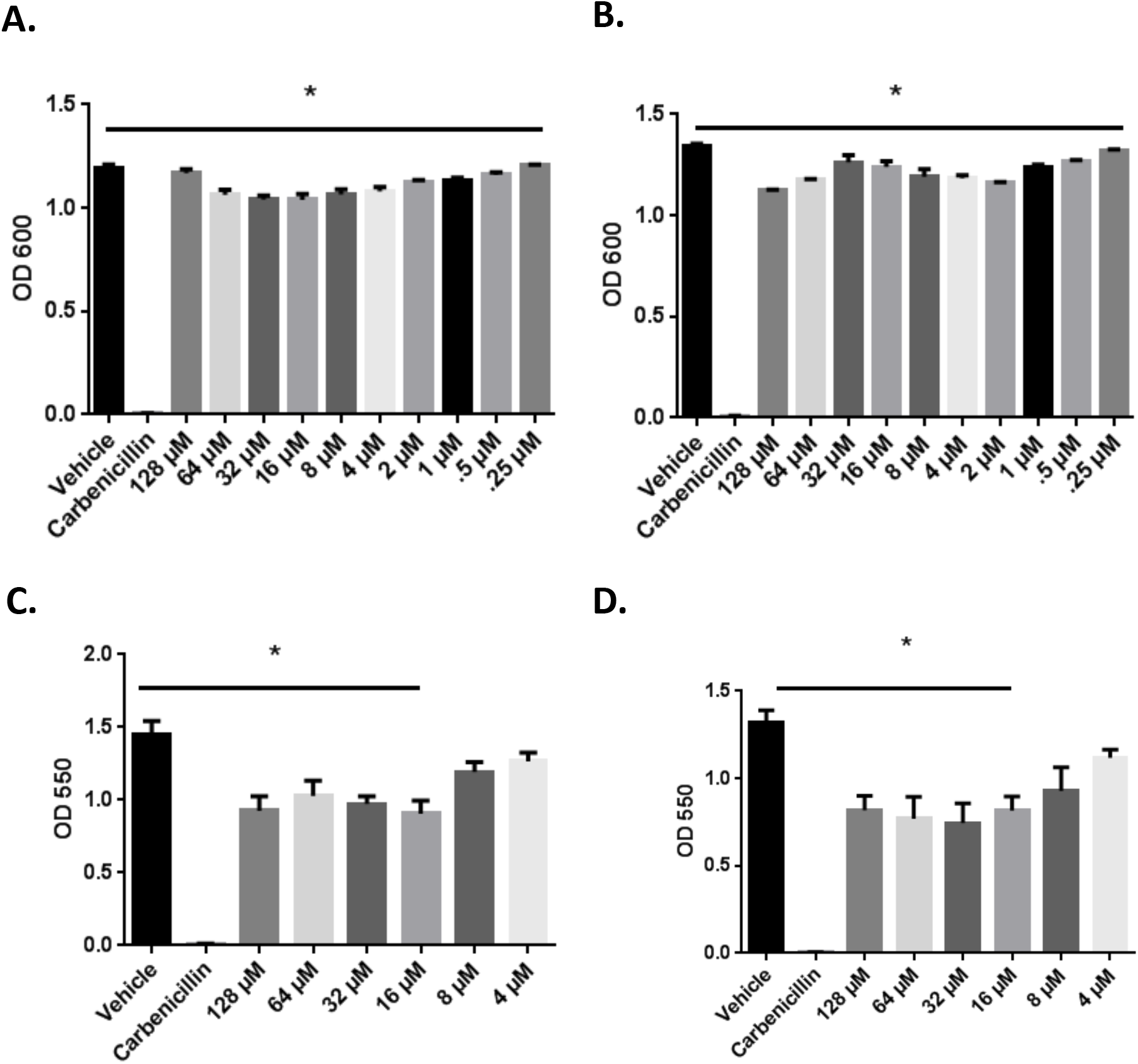
Antibiofilm activities of peptides 11 and 16 on *P. aeruginosa*. **A.** Peptide 11 and **B.** peptide 16 exhibit no bacteriostatic activity. **C.** Peptide 11 exhibits mild antibiofilm activities **D.** Peptide 16 exhibits mild antibiofilm activity. Data is representative of 3 biological replicates. A * represents a p-value < .05 as determined via one-way ANOVA **(A,B)**. A * represents a significant difference as determined via Tukey HSD compared to the vehicle control **(C,D)**.

**Supplemental Table 1:** Cytotoxicity of 20-mer bacteriocins at 128 μM. Y indicates an increase in hemolysis or cytotoxicity at 128 μM. N indicates no increase in hemolysis or cytotoxicity at 128 μM.

| Peptide # | Charge | Helicity | Hydrophobic Moment | Cytotoxic? | Hemolytic? |
| --- | --- | --- | --- | --- | --- |
| 1 | 1 | 18.93 | 322.53 | N | N |
| 2 | 1 | 18.13 | 222.87 | N | Y |
| 3 | 7 | 22.27 | 835.73 | N | N |
| 4 | 7 | 22.41 | 758.92 | N | N |
| 5 | 7 | 22.7 | 599.8 | N | N |
| 6 | 8 | 22.43 | 503.81 | N | N |
| 7 | 7 | 22.11 | 566.58 | N | N |
| 8 | 6 | 22.94 | 611.8 | N | N |
| 9 | 5 | 22.25 | 467.52 | N | N |
| 10 | 6 | 22.16 | 466.17 | N | Y |
| 11 | 4 | 20.86 | 912.76 | N | N |
| 12 | 6 | 21.3 | 673.77 | N | N |
| 13 | 5 | 20.53 | 680.63 | N | N |
| 14 | 9 | 20.81 | 567.158 | N | N |
| 15 | 9 | 20.19 | 357.31 | N | N |
| 16 | 7 | 19.51 | 471.63 | N | N |

